# The *Zymoseptoria tritici* effector Zt-11 contributes to aggressiveness in wheat

**DOI:** 10.1101/2024.09.04.611169

**Authors:** Sujit Jung Karki, Paola Pilo, Colleen Lawless, Nikolaos Mastrodimos, Anna M. M. Tiley, James Burke, Angela Feechan

## Abstract

*Zymoseptoria tritici* is an ascomycete fungus and the causal agent of Septoria tritici leaf blotch (STB) in wheat. *Z. tritici* secretes an array of effector proteins that are likely to facilitate host infection, colonisation and pycnidia production. In this study we demonstrate a role for Zt-11 as a *Z. tritici* effector during disease progression. *Zt-11* is upregulated during the transition of the pathogen from the biotrophic to necrotrophic phase of wheat infection. Deletion of *Zt-11* delayed disease development in wheat, reducing the number and size of pycnidia, as well as the number of macropycnidiospores produced by *Z. tritici*. This delayed disease development by the Δ*Zt-11* mutants was accompanied by a lower induction of *PR* genes in wheat, when compared to infection with wildtype *Z. tritici*. Overall, these data suggest that Zt-11 plays a role in *Z. tritici* aggressiveness and STB disease progression possibly via a salicylic acid associated pathway.

## INTRODUCTION

The fungal pathogen *Zymoseptoria tritici* is the causal agent of Septoria tritici blotch (STB) disease in wheat, which causes on average, yield losses of around 20% on susceptible wheat cultivars in North-Western Europe (Fones and Gurr, 2015). Due to the rapid emergence of fungicide tolerance and evolution of virulence against resistant wheat cultivars, *Z. tritici* poses a threat to wheat production worldwide (Fones et al., 2020).

Successful plant colonization by filamentous plant pathogens is dependent on the ability of the pathogen to overcome host defences. This manipulation of host defence is achieved through the secretion of a repertoire of effector proteins (Dou and Zhou, 2012; Van der Does and Rep, 2017). These effector proteins are typically small, secreted proteins (SSPs) that facilitate suppression of host cellular defence and/or protection from antimicrobial proteins (Presti and Kahmann, 2017). *Z. tritici* secretes SSPs that play a role in virulence and aggressiveness (Amaral et al., 2012; Gohari et al., 2015; Rudd et al., 2015; Karki et al., 2021). For example, two lysin motif containing effectors; Mg3LysM and Mg1LysM play an important role in protection of the fungal cell wall of *Z. tritici* against host chitinases through chitin sequestration (Marshall et al., 2011; Sanchez-vallet et al., 2020). Zt80707 is a secreted protein under positive selection that contributes to *Z. tritici* virulence (Poppe et al., 2015). Previously we found a conserved effector ZtSSP2, that can interact with a wheat host E3 ubiquitin ligase (TaE3UBQ) to promote STB infection (Karki et al., 2021).

Plants have also evolved receptors that can recognise some of these effectors and thus confer resistance to invading pathogens (Jones and Dangl, 2006; De Wit et al., 2009). Studies utilising association genetic mapping identified the *Z. tritici* effector gene *AvrStb6* (Zhong et al., 2017). The *Z. tritici* effector AvrStb6 is recognised by wheat cultivars carrying the corresponding resistance gene, *Stb6* which encodes a wall associated kinase (Saintenac et al., 2018). The recently cloned resistance gene *Stb16q* provides broad spectrum resistance and encodes a cysteine rich receptor-like kinase (CRK) which recognises the avirulence gene product AvrStb16q (Saintenac et al., 2021). While Stb15 which recognises AvrStb15, is a lectin receptor-like kinase (Hafeez et al., 2023). Finally, Avr3D1 is recognised by wheat cultivars carrying the resistance gene Stb7 (Meile et al., 2018)

In this study, we show a role for the *Z. tritici* effector protein; Zt-11 in disease development. Zt-11 is a small (79 amino acid), cysteine rich (10 cysteine residues) secreted protein located on core chromosome 5 (Gohari et al., 2015). It was previously found to interact with the wheat secreted proteins TaSRTRG6 (*Triticum aestivum* Septoria Responsive Taxonomically Restricted Gene 6) (Brennan et al., 2020), TaSSP6 and TaSSP7 (Zhou et al., 2020). Zt-11 appears to be specific to *Z*.

*tritici* and since it interacts with several host defence proteins, one of which (TaSRTG6) is taxonomically restricted, we studied *Zt-11* further to investigate a potential role in the *Z. tritici-* wheat interaction. *Zt-11* expression peaks at the transition from the biotrophic to the necrotrophic phase of STB infection. Using a targeted gene replacement approach to knockout *Zt-11* we demonstrate its role in the timing of aggressiveness, disease development and asexual sporulation of *Z. tritici* which is associated with salicylic acid (SA) related gene expression.

## MATERIALS AND METHODS

### Plant Material and Fungal Strains

The susceptible wheat cultivar Longbow was used throughout this study. Plastic trays containing John Innes Compost No. 2 (Westland Horticulture, UK) were sown with 10-12 wheat seeds. The wheat plants were grown in a growth chamber at 16 hours day/8-hour night photoperiod at 13,000 lux, RH 80% ± 5% at 19°C/20°C.

Cultures of *Z. tritici* IPO323 (Kema and Van silfhout, 1997) were grown on Yeast Peptone Dextrose Agar (YPDA) (yeast extract 10 g/l, peptone 20 g/l, glucose 20 g/L and technical agar 20 g/L) or Potato Dextrose Agar (PDA) (potato dextrose broth 24 g/L, technical agar 20 g/L) for five to six days in a growth cabinet with a temperature of 20 °C and 16:8 light/dark cycles. For *in vitro* phenotyping fungal strains were grown in Yeast extract glucose agar (YEG) (1% Yeast extract, 2% glucose and 2% Agar) medium supplemented with H_2_O_2_ at concentrations 0, 2, 4 and 6mM and the plates were incubated for 7 days at 20°C.

### Validation of protein secretion using a yeast sucrose secretion system

A Gateway-compatible vector (pGADT7) for yeast secretion assay and suc2 yeast mutant (strain SEY6210) was utilized (Brennan et al., 2020; Zhou et al., 2020). Briefly, the invertase (*SUC2*) gene with and without signal peptide was amplified from the yeast strain BY4741 with a linker (Kex2 site) added between the Gateway reading frame and the *SUC2* gene. This construct was ligated into the pGADT7 vector and verified by sequencing. Candidate *Zt-11* and Δ*SP-Zt-11* were cloned into the yeast secretion vector in-frame with the N-terminus of the SUC gene and transformed into the suc2 yeast mutant. Transformants were PCR validated using primers listed on Supplementary Table 1 and selected on a synthetic dropout medium (minus Trp and Leu) with sucrose as a sole carbon source. Yeast spotting was performed with dilutions of 10^−1^, 10^−2^, and 10^−3^ respectively. The experiment was repeated twice independently with four replicates per independent experiment.

### RNA extraction and quantitative RT-PCR

For the *Zt-11* gene expression studies, 14 day-old wheat leaves infected with *Z. tritici* IPO323 were collected at 0, 4, 8, 10, 13, 15, 17 and 21 days post-infection (dpi). Each sample consisted of 100mg pooled from two individual leaves from each seedling. Samples were immediately frozen in liquid nitrogen and stored at –80 °C. The experiment was repeated three times independently.

For plant defence gene expression, 14 days old wheat leaves were infected with either wild type strain (IPO323) or KO strains (designated Δ*Zt11_2B,* Δ*Zt11_3E* and Δ*Zt11_7A*). Sampling was carried out at 0, 2, 6 and 10 days post inoculation (dpi). All samples (100mg) were immediately frozen in liquid nitrogen and stored at −80°C until use. Defense marker genes were analyzed by real-time quantitative reverse transcription PCR (qRT-PCR) with primers for the following: pathogenesis-related (*PR*) genes *TaPR1*, *TaPR2*, and *TaPR5*; superoxide dismutase (*TaSOD*) and catalase (*TaCAT*). Primers are listed in Supplementary Table 1. Two independent experiments were carried out. Each experiment included three leaves each from three plants per strain per time point (*n* = 6).

Total RNA was extracted from *Z. tritici*-infected wheat leaves or YPDA cultures using the RNeasy Mini Kit (Qiagen) following the manufacturer’s instructions. RNA was then subjected to on-column DNase treatment (Sigma). Quantification of total RNA was carried out using a Nanodrop ND-1000 spectrophotometer. Reverse transcription of 1–2 μg of RNA for cDNA synthesis was carried out using the Omniscript RT Kit (Qiagen).

Real-time quantitative PCR (qRT-PCR) was carried out in 12.5 μl reactions including 1.25 μl of a 1:5 (v/v) dilution of cDNA, 0.2 μM of primers, and 1×SYBR Premix Ex Taq (Tli RNase H plus, RR420A; Takara). PCR conditions were as follows: 1 cycle of 1 min at 95 °C; 40 cycles of 5 s at 95 °C and 20 s at 60 °C; and a final cycle of 1 min at 95 °C, 30 s at 55 °C, and 30 s at 95 °C for the dissociation curve. qPCR was performed using the QuantStudio 7 Flex Real-Time PCR system (Applied Biosystems) and the relative gene expression was calculated as 2^−(Ct target gene–Ct housekeeping gene) as previously described (Livak and Schmittgen, 2001). The *Z. tritici* L-tubulin gene (Rudd et al., 2015) was used as reference gene for the *Zt-11* time course. Wheat *glyceraldehyde 3-phosphate dehydrogenase* (*TaGAPDH*) was used as a reference gene for defence gene quantification. Average Ct and SEM values were calculated from six individual Ct values per strain per time point.

### Construct Generation

The gene deletion construct for *Zt-11* was constructed using a yeast-based homologous recombination as described by Tiley et al. 2019. Primers for plasmid construction are listed in Supplementary Table 1. Briefly, the 1.5 kb upstream and downstream flanking region was selected for the effector gene candidate *Zt-11* (*Mycgr3G104444*), based on the reference genome database available for *Z. tritici* IPO323 available at JGI (Goodwin et al., 2011). Yeast based homologous recombination was utilized for knockout (KO) plasmid construct using the pCAMBIA0380_YA (yeast-adapted) vector as a backbone, Hygromycin-*trpC* resistance cassette from pCB1003 (Carroll et al., 1994) flanked by two 1.5 kb regions targeting the *Zt-11* locus. The flanking regions and Hygromycin-*trpC* resistance cassette were amplified using Phusion® High-Fidelity DNA Polymerase (Thermo Fisher Scientific) and successive PCR product purification.

A Zymoprep™ Yeast Plasmid Miniprep II kit (Zymo Research) was used to recover plasmid DNA from *Saccharomyces cerevisiae*, propagated into *Escherichia coli* DH5α cells and isolated using the GeneJET Plasmid Miniprep Kit (Thermo Fisher Scientific) or Gene JET Plasmid Midiprep Kit (Thermo Fisher Scientific) following the manufacturer’s instructions. Correct plasmid assembly was confirmed by PCR and sequencing.

### *Agrobacterium*-Mediated Transformation

The Δ*Zt-11* knockout vector was transformed into *Agrobacterium tumefaciens* LBA1126 and AGL1 cells. Agrobacterium–mediated transformation was then used to transform the *Z. tritici* IPO323 strain following the protocol outlined in Derbyshire et al., (2015; 2018).

### Confirmation of Δ*Zt-11* strains

Following 10-14 days after transformation separate *Z. tritici* transformants were selected and transferred to YPDA (10 g/L yeast extract, 20 g/L peptone, 20 g/L glucose, and 20 g/L technical agar) supplemented with Hygromycin B (100 μg/ml) and Timentin^TM^ (100 μg/ml). Potential mutants were then sub-cultured at least three times to acquire a pure culture. Double PCR was carried out on the *Z. tritici* genomic DNA extracted with the sonication method (Pilo et al., 2022) using two primer pairs; the first primer pair (Fwd_Zt11_WT and Rvs_Zt11_WT) was designed to amplify the wild-type gene, and the second pair (Fwd_Hyg and Rvs_Hyg) to amplify the Hygromycin-trpC resistance cassette (Supplementary Figure 1). We further confirmed the deletion of *Zt-11* gene by sequencing the insertions region (flanking regions and hygromycin insert) on all three independent KO strains (Δ*Zt11_2B,* Δ*Zt11_3E* and Δ*Zt11_7A*) (Supplementary Figure 2). Complementation of the mutants was unsuccessful; thus, an ectopic strain was selected in order to validate the mutants. In the ectopic strain both bands are present due to the insertion of the Hygromycin-trpC resistance cassette in other genome coordinates with no deletion of the *Zt-11* gene. Primers used for knockout confirmation and sequencing are listed in Supplementary Table 1.

### *In planta* Experiments

For plant infections, the *Z. tritici* IPO323, ectopic and three independent KO strains (designated Δ*Zt11_2B,* Δ*Zt11_3E* and Δ*Zt11_7A*) were cultured onto YPDA and grown at 20°C under white light supplemented with blue/black ultraviolet (UV-A) light under a 12:12 hour light:dark photo cycle for approximately 7 days to achieve yeast-like growth of fungal isolates (Tiley et al., 2018). Fungal spores from the YPDA cultures were carefully harvested using an inoculating T-shaped spreader and suspended in deionised sterile water. Spore concentrations were adjusted to 1x10L ml^-1^ in deionised sterile water containing 0.01% Tween20 solution. A spore suspension of 20 ml per plant tray was sprayed using plastic hand-held spray bottles. Control plants were sprayed with 20 ml of deionised sterile water containing 0.01% Tween20. The first true leaf of the wheat cv. Longbow was held adaxial side up on polystyrene blocks for inoculation (Keon et al., 2007). Wheat seedlings were inoculated at 14 days old. At least 10-12 seedlings were potted in each tray for each strain per independent experiment. Inoculated plants were then covered with polythene bags for 72 h to ensure high humidity. Four independent experiments were carried out n ≥ 40 plants per strain.

Virulence and aggressiveness of *Z. tritici* Δ*Zt-11 strains*, IPO323 and the ectopic strain was assessed by monitoring and recording disease progression from the appearance of the first symptoms at 8 dpi. Disease symptoms on the infected leaves were scored from 1 to 5 by eye using a modified version of the scale from Skinner (2001). Briefly, the scale measurement was as follows: 1 representing no disease symptoms, 2 representing small chlorotic flecks, 3 representing chlorosis, 4 representing necrosis and lastly 5 representing necrosis with pycnidia.

### Pycnidia and pycnidiospore production

Pycnidia production was analysed for each strain at 21 dpi. A total of ten leaves were sampled from each strain for four independent experiments (n = 40 per strain). Leaves were scanned (1200 dots per inch) using an Epson Perfection V600 Photo Scanner and analysed by ImageJ (Schindelin et al., 2015) with an automated technique protocol (Stewart et al., 2016; Karisto et al., 2018). Each leaf was designated a QR code and following parameters were automatically recorded from the scanned image: total leaf area, necrotic and chlorotic leaf area, number of pycnidia, and their positions on the leaf. Pycnidia production per centimetre square of leaf was calculated for the IPO323, ectopic, and Δ*Zt-11* strains.

The same leaves which were used for pycnidia production were then used to calculate the number of pycnidiospores. Six centimetres from each leaf, for a total of ten leaves for each strain (n = 40), were placed in a petri dish in high humidity for 48h at room temperature (20 ± 3°C). Each leaf was then suspended in 2 ml of sterile distilled water and vortexed for 5 seconds. Suspended macro pycnidiospores and micropycnidiospores were counted with a haemocytometer per each leaf and recorded (Tiley et al., 2019). Four independent experiments were carried out.

For pycnidia size measurements, two leaves from two plants per isolate at 21 dpi were harvested and stained with trypan blue. The leaf sections were then mounted with Polyvinyl Alcohol-Lactic acid (PVA-L) medium (8% v/v) microscopy glue and observed under light microscope (Leica DM5500B). Pycnidia width was calculated using the custom scale bar in Leica DM5500B microscope. This experiment was conducted twice independently with at least 100 pycnidia counted per leaf per plant.

## RESULTS

### Zt-11 is a small, secreted protein specific to *Z. tritici*

Eight *Z. tritici* SSPs were previously shown to interact with wheat proteins that play a role in defence against *Z. tritici* (Brennan et al., 2020; Zhou et al., 2020). We conducted a BLAST_P_ search to identify any potential homologues of these eight *Z. tritici* SSPs (Supplementary Table 2). Two SSPs from *Z. tritici* (Zt-11 and Zt-26) out of the eight lacked hits in any fungal species except *Z. tritici* (Supplementary Table 2).

Zt-11 is predicted to have a N-terminal signal peptide (0.9978 likelihood, SignalP-5.0) (Almagro et al., 2019). The secretion signal was validated using a yeast secretion assay (Figure 1A; (Brennan et al., 2020; Zhou et al., 2020). The full length *Zt-11* construct was able to complement the *suc2* knockout yeast strain, thus allowing the yeast cells to grow in selection media containing sucrose as the sole source of carbon (Figure 1B). No conserved domains were found for Zt-11 using the NCBI Conserved Domain Database (NCBI CDD) (Marchler et al., 2015).

**FIGURE 1.**
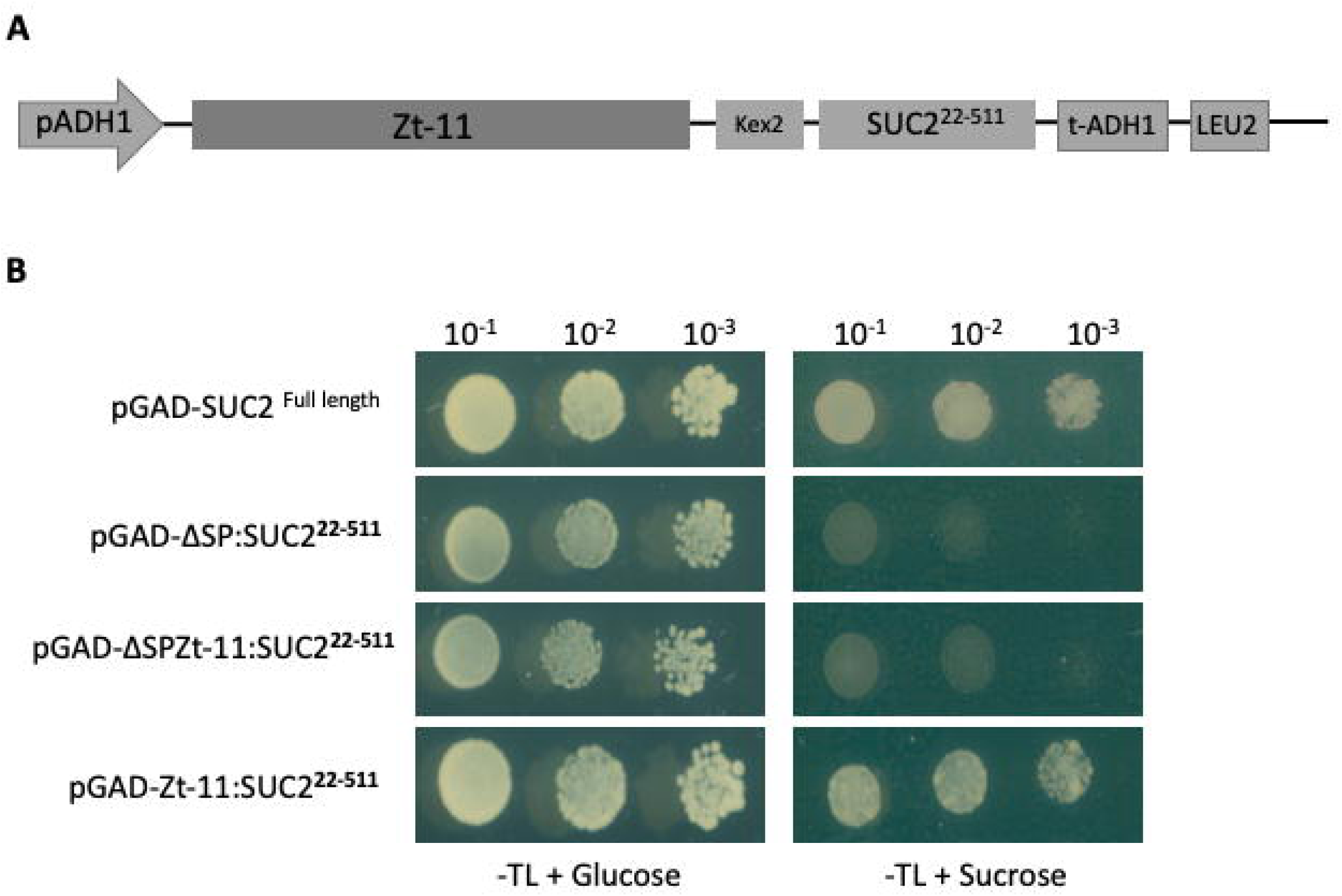
*Zt-11* is a secreted protein. (**A)** Design of the gateway compatible yeast pGADT7-Zt- 11-Suc2 vector and Invertase mutant yeast strain SEY6210 used for the secretion assay (Brennan et al. 2020). (**B)** Yeast strains carrying *Zt-11* with secretion signal fused in frame with the Invertase gene *Suc2* were able to grow in sucrose containing drop out media (SD-TL) sucrose therefore cells will grow if Invertase is secreted. SEY6210 carrying the pGAD-ΔSP:SUC2^22-511^ vector was used as negative control while SEY6210 with pGAD-SUC2^Full^ ^length^ acts as positive control. This experiment was repeated twice independently with four replicates per construct in each experiment.

The timing of *Zt-11* expression during infection of wheat was investigated using reverse transcription-quantitative polymerase chain reaction (RT-qPCR). During wheat infection, *Zt-11* was significantly upregulated (*P* < 0.01) from 4 dpi to 10 dpi compared to 0 dpi, with a peak of expression at 10 dpi. At the later necrotrophic phase of infection (15-21 dpi) *Zt-11* expression was downregulated with low expression levels similar to those found at 0 dpi. *Zt-11* was expressed at low levels *in vitro* compared to *in planta* conditions (Figure 2).

**FIGURE 2.**
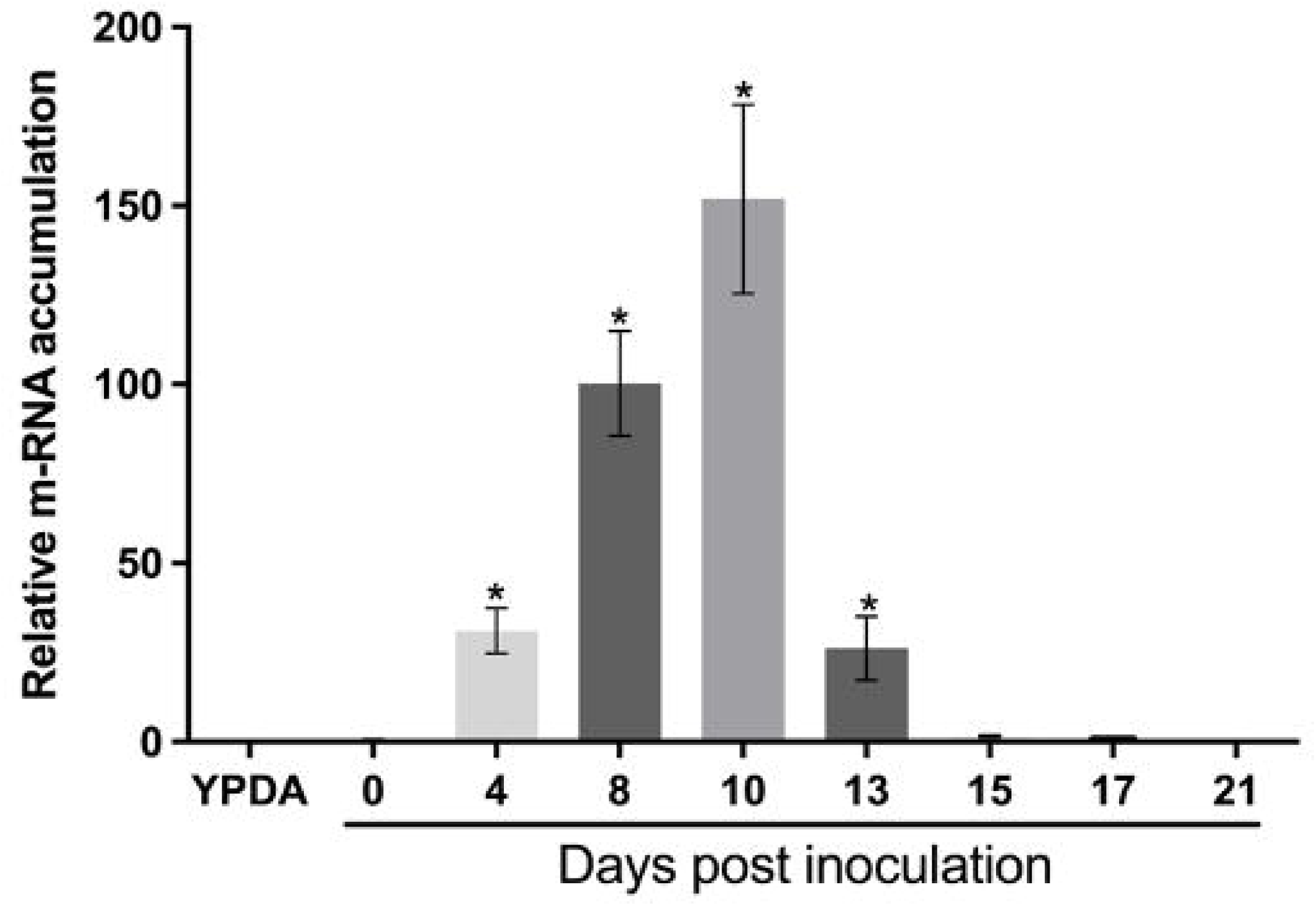
*Zt-11* expression during *Z. tritici* infection. Transcript levels of *Zt-11* during *in vitro* (YPDA) compared with *in planta* infection of wheat at 0, 4, 8, 10, 13, 15, 17 and 21 days post inoculation (dpi) (cv. Longbow) with *Z. tritici* (IPO323). The expression levels of β*-tubulin* (*Z. tritici*) were used to normalize the expression levels of *Zt-11*. Each independent experiment had two leaves each from two individual plants. Three independent experiments were performed. The bars represent the mean relative expression ± SEM. Asterisk above bars indicate significant differences compared to 0 dpi, as determined by Tukey’s test (*P* < 0.01).

### Δ*Zt-11* strains show delayed disease development

To understand whether Zt-11 plays a role in *Z. tritici* virulence and aggressiveness, we generated *Zt-11* deletion strains using *Agrobacterium tumefaciens*-mediated transformation (Supplementary Figure 1A). Successful transformation was indicated by loss of the wild-type *Zt-11* and the gain of the knockout (KO) fragment (Supplementary Figure 1B). Additionally, *Zt-11* deletion strains were also confirmed by amplicon sequencing (Supplementary Figure 2). Three independent deletion strains (Δ*Zt-11_2B,* Δ*Zt-11_3E* and Δ*Zt-11_7A*) and an ectopic strain were selected.

The parental IPO323 wild-type (*WT*) strain, the KO strains Δ*Zt-11_2B,* Δ*Zt-11_3E* and Δ*Zt-11_7A* and ectopic strain were subsequently inoculated onto leaves of the susceptible wheat cultivar Longbow. Disease progression was monitored from 3 dpi and recorded from 8 dpi when the first signs of disease symptoms were observed. The first symptoms of disease were visible as small chlorotic lesions observed on leaves infected with the IPO323 and ectopic strain at 10 dpi (Figure 3A). The chlorotic lesions developed into necrotic lesions with pycnidia, visible from 13-15 dpi on leaves infected with IPO323 and the ectopic strain (Figure 3A). However, the leaves infected with Δ*Zt-11* strains developed chlorotic lesions only from 13 dpi, followed by necrosis and pycnidia production from 17 dpi (Figure 3A). Overall, the timing of symptom appearance for the Δ*Zt-11* strains was significantly delayed (*P* < 0.01) with lower disease scores at 10, 13 and 15 dpi compared to IPO323 and the ectopic strain (Figure 3B).

**FIGURE 3.**
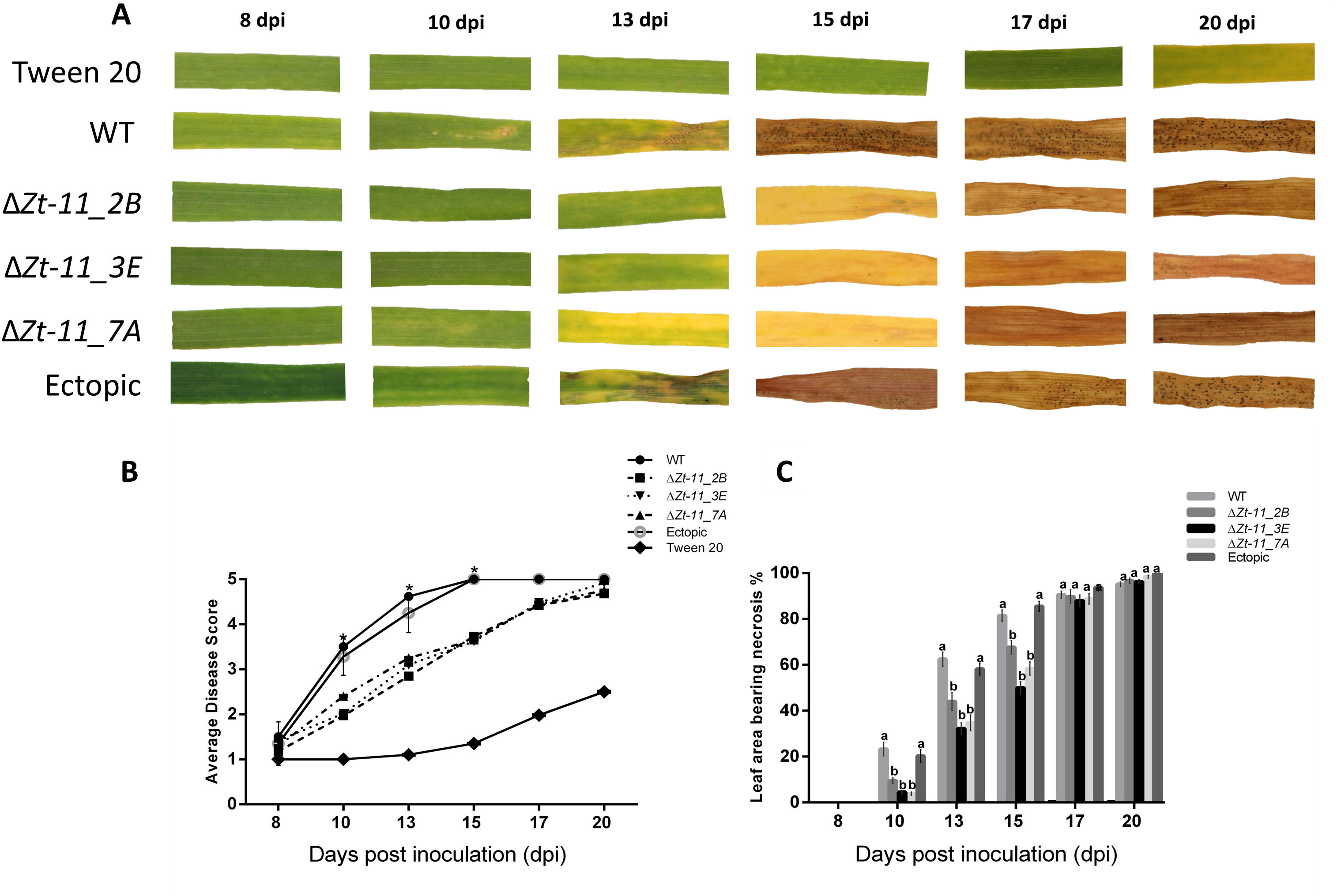
Δ*Zt-11* mutant strains show delayed disease development. (**A)** Disease development of Δ*Zt-11* mutant on the susceptible wheat cv. Longbow compared to IPO323 and ectopic strain. Images are representative of four independent experiments (n ≥ 40). At least 10 leaves each from an individual plant per experiment were analysed. (B) Disease progression of wheat infected with *Z. tritici* Δ*Zt-11* mutant strains compared with IPO323 and the ectopic strain from 8 dpi to 20 dpi based on the 1 to 5 scale (Skinner, 2001). Each data point represents the average of four independent experiments, with at least 10 leaves per experiment per strain, each from an individual plant (n ≥ 40). Asterisks represent significant differences between the IPO323 strain and mutants (ΔZ*t-11_2B*, Δ*Zt- 11_3E* and Δ*Zt-11_7A*) at P < 0.01, according to Mann-Whitney-U-test test. (C) Necrosis percentage of wheat cv. Longbow leaves infected with Δ*Zt-11* mutant strains compared with the IPO323 and ectopic strain from 8 dpi to 20 dpi. Bar chart showing necrosis progression with each data point represents the average of four independent experiments with at least 10 leaves each from an individual plant per experiment per isolate (n ≥ 40). Error bars are +/- SEM of the mean. Different letters indicate significant differences between each other (Tukey’s HSD test, P < 0.01).

To further investigate the delayed disease symptoms displayed by leaves infected with the Δ*Zt-11* strains, we used percentage leaf area bearing necrosis as an additional disease index (Figure 3C). A significant reduction (*P* < 0.01) in the percentage of leaf area covered in necrosis was observed for the Δ*Zt-11* strains at 10, 13 and 15 dpi when compared to IPO323 and the ectopic strain (Figure 3C). However, from 17 dpi onwards there were no significant differences in necrosis between Δ*Zt- 11* strains with the ectopic strain as well as IPO323.

### Deletion of *Zt-11* impacts asexual sporulation

Further analysis was performed to determine if *Zt-11* impacts pycnidia production. We measured pycnidia density on infected leaves (Karisto et al., 2018). At 21 dpi, the number of pycnidia on each leaf of the Δ*Zt-11* strains was determined and compared to the IPO323 strain (Figure 4A). The Δ*Zt- 11* strains produced a significantly smaller number (*P* < 0.01) of pycnidia, in comparison to IPO323 and the ectopic strain. On average the mutants produced significantly less pycnidia compared to leaves infected with IPO323 and the ectopic strain (Figure 4A). In addition to differences in the number of pycnidia produced by the Δ*Zt-11* strains, we also measured the pycnidia size (Figure 4B, C) and found that Δ*Zt-11* strains produced significantly smaller pycnidia (*P* < 0.01) compared to the IPO323 and ectopic strain (Figure 4B, C). We found no significant differences in pycnidia production between the ectopic strain and IPO323.

**FIGURE 4.**
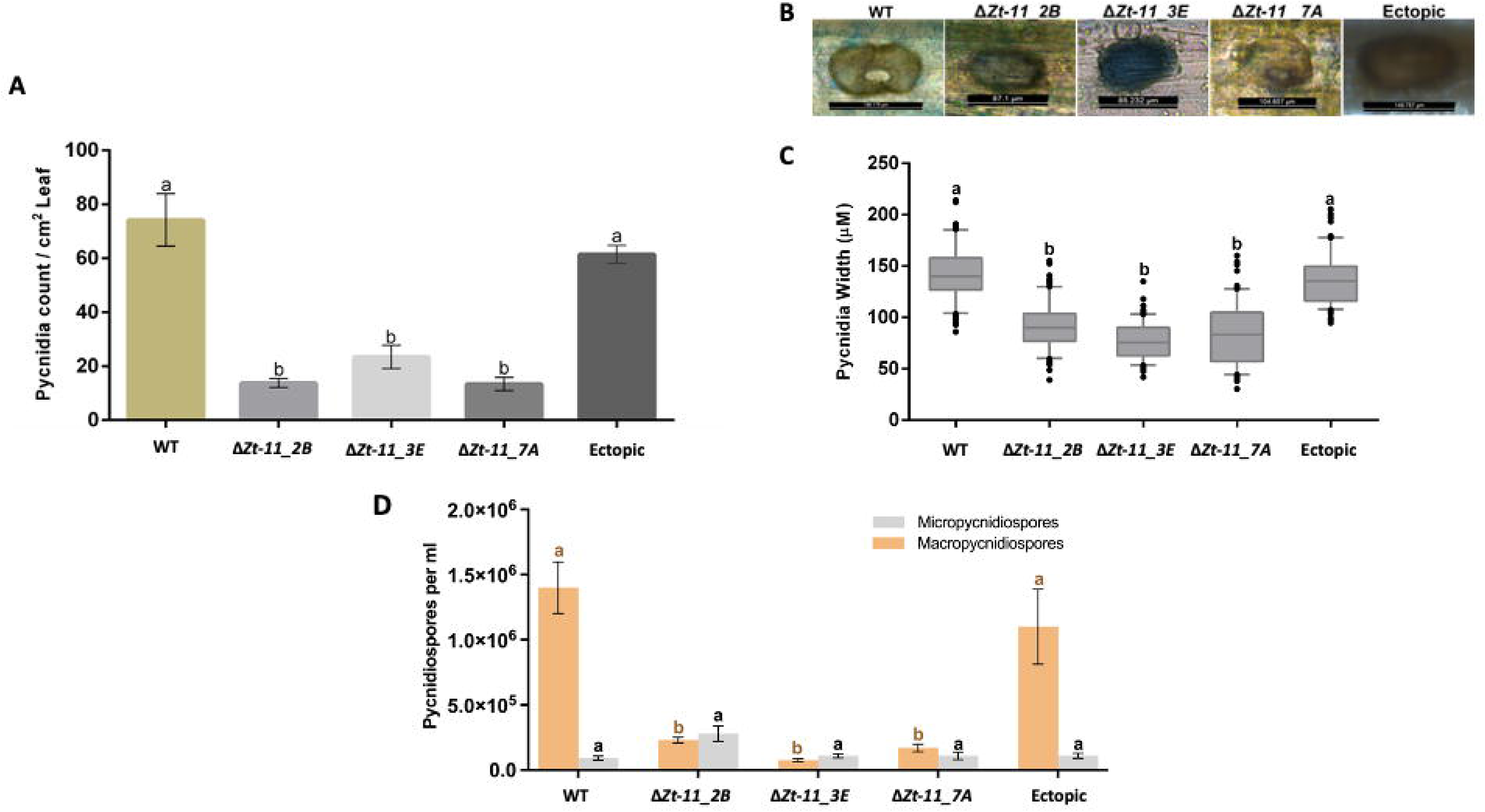
Deletion of *Zt-11* impacts asexual sporulation. (**A)** Δ*Zt-11* mutants produced significantly fewer pycnidia compared to the IPO323 and ectopic strain. Inoculated leaves were scanned at 21 dpi and scanned images were analysed to obtain the number of pycnidia with the software Image J. Each data point represents the average of four independent experiments with at least 10 leaves per experiment per strain, each from an individual plant (n ≥ 40). (**B)** Representative images of Pycnidia size between the IPO323, ectopic and Δ*Zt-11* mutant strains. (**C)** Comparison of pycnidia size between IPO323, ectopic and *Zt-11* mutant strains. For pycnidia size measurement, two leaves from two plants per isolate at 21 dpi were taken and stained with trypan blue and at least 100 pycnidia were counted per leaf using a light microscope (Leica DM5500B). Pycnidia width was calculated using the scale bar. The experiment was conducted twice independently. Error bars are +/- SEM of the mean. Different letters indicate significant differences between each other (Two- way ANOVA and Tukey’s HSD test, *P* < 0.01). (D) Stacked bar chart showing macropycnidiospores and micropycnidiospores counts per ml at 21 dpi (Leaf= 6cm). Δ*Zt-11* mutants produced significantly lower macropycnidiospores numbers (indicated by different letters) but not fewer micropycnidiospores when compared to the IPO323 and ectopic strain. Each data point represents the average of four independent experiments, 10 leaves per experiment per strain, each from an individual plant (n ≥ 40). Different letters indicate significant differences between each other (Two-way ANOVA and Tukey’s HSD test, P < 0.01).

The Δ*Zt-11* strains were able to exude cirrhus containing pycnidiospores (Supplementary Figure 3). The asexual spores produced by *Z. tritici* fall into two categories: macropycnidiospores, and micropycnidiospores, (Eyal et al., 1987: Tiley et al., 2019). Since *Zt-11* impacted pycnidia number and size we assessed the types of pycnidiospores produced. The number of macropycnidiospores and micropycnidiospores in the Δ*Zt-11* strains were counted and compared to IPO323 and the ectopic strain (Figure 4D). Macropycnidiospores production was significantly lower (80 ± 10%, *P* < 0.01) in leaves infected with the Δ*Zt-11* strains compared to IPO323 and the ectopic strain (Figure 4D). While, the number of micropycnidiospores produced was not significantly different between strains (Figure 4D).

### Δ*Zt-11* strains lead to lower induction of host SA mediated defence genes during infection

To further determine whether delayed disease development by the mutant strains were due to changes in host defense gene related transcripts, qRT-PCR was used to examine the levels of wheat pathogenesis-related genes namely; *TaPR1*, *TaPR2* and *TaPR5* as markers of systemic acquired resistance (SAR) in plants (Figure 5A-C) (Shah and Zeier, 2013; Ajigboye et al., 2021; Li et al., 2023) and two reactive oxygen species (ROS) related wheat genes Catalase (*TaCAT)* and Superoxide dismutase (*TaSOD*) (Figure 5D, E). The wheat plants infected with the *Z. tritici* strain IPO323 showed early induction of *TaPR1* and *TaPR5* at 2 dpi and also had significantly higher transcript accumulation of *TaPR1* at 10 dpi compared to 0 dpi. The levels of *TaPR1, TaPR2* and *TaPR5* were significantly lower in wheat infected with the Δ*Zt-11* mutant strains than IPO323 at all timepoints with the exception of *TaPR2* at 0 dpi and 10 dpi and *TaPR1* at 6 dpi (Figure 5A-C). The levels of *TaCAT* were significantly upregulated at 10 dpi compared to 0, 2 and 6 dpi for all *Z. tritici* strains. *TaSOD* was downregulated at 6 dpi compared to 0, 2 and 10 dpi for all *Z. tritici* strains. However, the transcript level of ROS scavenging genes *TaSOD* and *TaCAT* were not significantly different in infected wheat leaves between IPO323 or Δ*Zt-11* mutant strains (Figure 5D, E). We investigated the vegetative growth of the Δ*Zt-11* mutants on solid media (PDA and YEG) to determine the impact of H_2_O_2_ (2, 4 and 6 mM) on the Δ*Zt-11* mutants compared to IPO323. In line with no differences in antioxidant (*TaSOD* and *TaCAT*) gene expression between IPO323 and the Δ*Zt-11* strains, we observed no differences in vegetative growth, melanisation, nor in resistance/susceptibility to H_2_O_2_ between IPO323 and Δ*Zt-11* mutants (Supplementary Figure 4).

**FIGURE 5.**
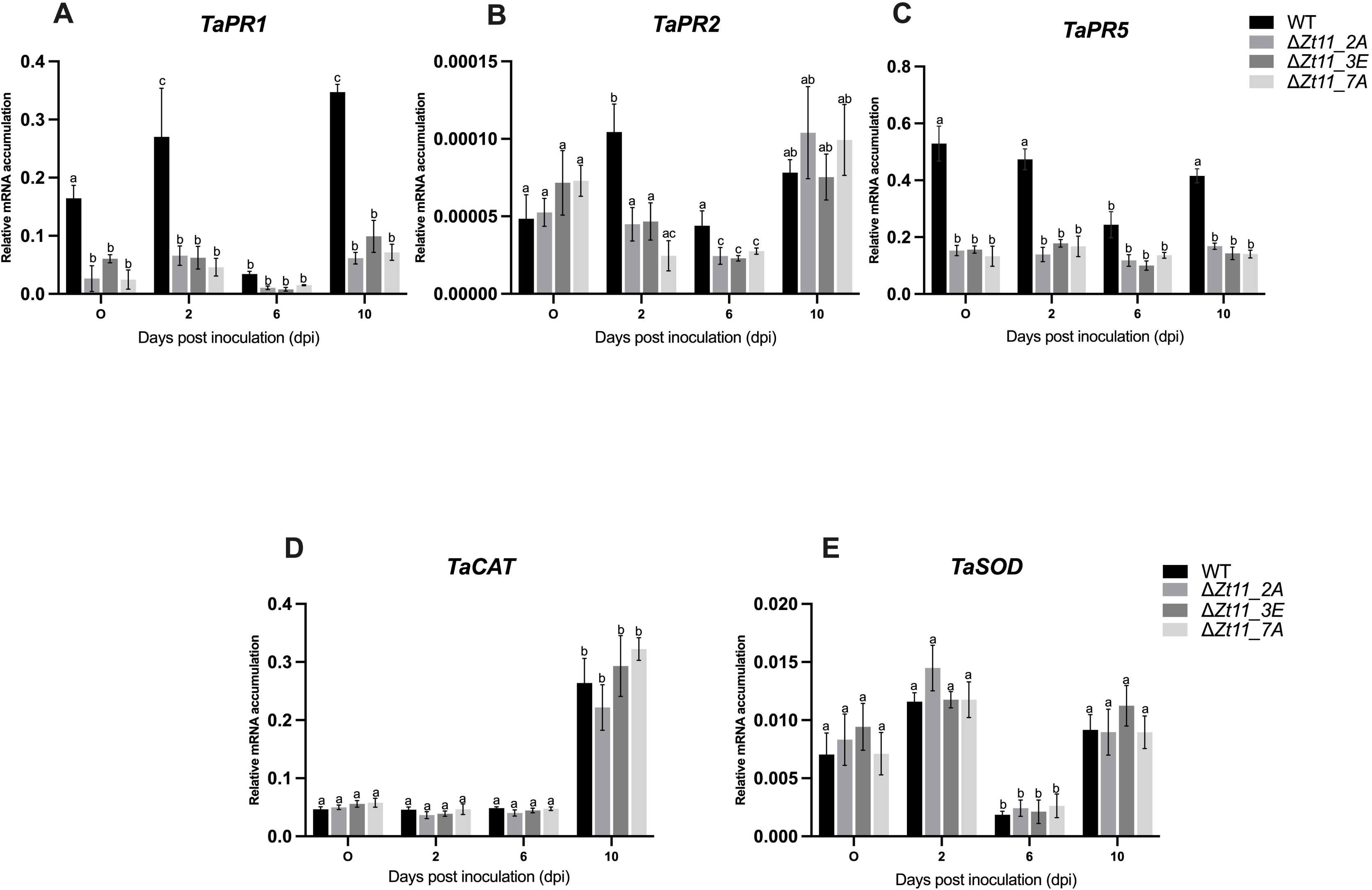
Transcript profile of wheat defence genes in wheat samples infected with the Δ*Zt-11* mutant strain compared with IPO323. (A) *TaPR1,* (B)*TaPR2*, (C) *TaPR5*, (D) *TaCAT* and (E) *TaSOD*. The results represent the mean of three leaves (each from individual plants) per strain per time point over two independent experiments (*n* = 6) (error bars indicate ±SEM). Relative gene quantification was calculated by the comparative 2^-ΔCT method (Livak and Schmittgen, 2001). Normalization was carried out using wheat *glyceraldehyde 3-phosphate dehydrogenase*. Different letters indicate significant differences between each other as determined using Tukey’s HSD test. (P < 0.01).

## DISCUSSION

*Z. tritici* is known to secrete an abundant number of effector proteins into the wheat host apoplast during infection (Gohari et al., 2015; Rudd et al., 2015; Kettles et al., 2017). Effectors are at the frontline during plant-pathogen interactions and a diverse array of these pathogen effectors are known to interact with host targets to allow infection and host colonization (Kamoun, 2006; Hogenhout et al., 2009; Rafiqi et al., 2013). Owing to the importance in manipulation of host processes, studies have highlighted that most pathogen effector genes are cysteine rich secreted proteins and have increased expression in host plants. This reflects their function compared to *in-vitro* conditions (Gohari et al., 2015; Rudd et al., 2015; Palma-Guerrero et al., 2017). *Zt-11* was predicted to be a secreted protein, based on the presence of N-terminus signal peptide sequence and this was validated by using a secretion assay (Brennan et al., 2020; Zhou et al., 2020).

*Z. tritici* has been described as hemibiotroph (Rudd et al., 2015) or a late necrotroph with a long latent phase (Sanchez-vallet et al., 2015). We examined the expression of *Zt-11* in infected wheat leaves using qRT-PCR and found that *Zt-11* displayed significantly induced expression *in planta* from 4 dpi to 10 dpi. This expression profile of *Zt-11* corresponds to the switch from the biotrophic to necrotrophic phase of *Z. tritici* infection. A similar expression pattern of *Zt-11* was also reported previously where *Zt-11* expression was low at 2 dpi peaking at 8 dpi before decreasing from 12-20 dpi (Gohari et al., 2015; Palma Guerrero et al., 2017).

Our previous studies indicated that Zt-11 interacts with wheat proteins TaSRTRG6, TaSSP6 and TaSSP7 (Brennan et al., 2020; Zhou et al., 2020). These are small, secreted wheat proteins highly expressed in response to *Z. tritici* infection. TaSRTRG6 has homology with a Bowman-Birk type proteinase inhibitor (Brennan et al., 2020). In rice, a similar proteinase inhibitor AvrPiz-t Interacting Protein 4 (APIP4) is an effector target protected by a nucleotide-binding and leucine-rich repeat (NLR) *R* protein (Piz-t) (Zhang et al., 2020). TaSSP6 has significant homology to a glycine rich protein (Zhou et al., 2020). Previously, a glycine rich protein in Lily (*Lilium* spp.) was found to be required for resistance against the necrotrophic pathogen *Botrytis elliptica* (Lin et al., 2020). TaSSP7 has significant homology to a papilin-like isoform which is an extracellular matrix glycoprotein (Kramerova et al., 2000). Silencing of these genes in wheat increased susceptibility to STB, confirming their role in defence against *Z. tritici* (Brennan et al., 2020; Zhou et al., 2020). Owing to its interaction with *Z. tritici* responsive wheat resistance protein, we hypothesize that *Zt- 11* could be an important effector protein of *Z. tritici* and plays a role in disease. Therefore, the contribution of *Zt-11* to the virulence and aggressiveness of *Z. tritici* was assessed using a targeted gene replacement approach to knockout *Zt-11*.

In *Z*. *tritici*, only a few small secreted protein effectors such as *LysM* and *Zt80707*, when deleted show an impact on fungal virulence (Marshall et al., 2011; Poppe et al., 2015). The *Z. tritici* mutants with deletion of *Zt-11* were able to infect wheat, however, the mutants were found to be significantly delayed in disease development from 10 dpi. This timing of significant lower disease symptoms coincides with the expression of *Zt-11* during the switch from the biotrophic to necrotrophic phase suggesting that *Zt-11* may support the timely progression to necrotrophy. It is possible that the slower wheat infection observed for the Δ*Zt-11* mutants compared to the wildtype *Z. tritici* strain is due to lower fungal biomass produced over time. *Z. tritici* growth rate can predict the latent period and pycnidia coverage (Rahman et al., 2020). We observed that Δ*Zt-11* strains were impaired in the development of pycnidia, producing significantly reduced numbers of pycnidia. The pycnidia of the Δ*Zt-11* strains were also significantly smaller in width compared with pycnidia from the IPO323 strain. This impaired pycnidia development may be a result of slower infection and lower fungal biomass produced by the Δ*Zt-11* mutants.

Previously, no significant difference in pathogenicity was found between a single Δ*Zt-11* strain and wildtype *Z. tritici*, following leaf infection with 1x10^7^spores/ml at 21 dpi (Mustafa et al., 2022). However, in this study significant differences in disease progression were found between three Δ*Zt- 11* strains and wildtype *Z. tritici* between 10-15 dpi using 1x10L spores/ml. The lower inoculum density coupled with a time course and three independent Δ*Zt-11* strains may explain the detection of significant differences in disease progression here compared to observation at 21 dpi only. In addition measuring pycnidia number using an automated technique coupled with microscopy led to the discovery of reduced pycnidia number and size as well as altered macropycnidiospore production *in planta*, all of which had not been previously assessed.

The asexual reproduction of *Z. tritici* comprises macropycnidiospore and micropycnidiospore production. Macropycnidiospores are curved and elongated around 35-98 µm x 1-3 µm in size with a total of 3-5 septa and micropycnidiospores are smaller 8-10.5 µm x 0.81 µm with no septa. Both spore forms can form hyphal growth, and both are able to infect wheat (Eyal et al., 1987; Perelló et al., 1990; Tiley et al., 2019). Our investigation of asexual spore types suggests that *Zt-11* deletion significantly decreases macropycnidiospores numbers rather than micropycnidiospores. Production of macropycnidiospores and micropycnidiospores are regulated by different pathways in *Z. tritici* (Tiley et al., 2019) and this suggests that *Zt-11* could contribute to macropycnidiospore production. The impaired pycnidia and pycnidiospores production of the Δ*Zt-11* strains in our study is similar with a study reporting the impact of the deletion of the extracellular effector, *Zt80707* in *Z. tritici* which also led to delayed pycnidia formation, lower pycnidia numbers, reduced pycnidiospores production and smaller pycnidia (Poppe et al., 2015).

Previously, a single Δ*Zt-11* strain was observed to have longer yeast-like spores (blastopores) produced *in vitro* (YPD) compared to wildtype *Z. tritici*. However, we did not observe morphological differences in yeast-like spores between the wildtype *Z. tritici* strain and three independently generated Δ*Zt-11* mutants *in vitro* (YPD). In addition, we did not find impaired mycelial growth across the three Δ*Zt-11* mutants compared to wildtype *Z. tritici* on PDA or YEG. In contrast, impaired mycelial growth was previously reported for a Δ*Zt-11* strain grown *in vitro* (PDA and water agar) (Mustafa et al., 2022).

Rapid induction of defence-related pathogenesis-related (*PR*) genes are triggered in wheat against infection by *Z. tritici* (Adhikari et al., 2007; Ray et al., 2003; Orton et al., 2017; Ors et al., 2018). We found that the expression of *PR* genes especially *TaPR1*and *TaPR5* were between 0 dpi and 10 dpi in Δ*Zt-11*-infected wheat samples compared to WT-infected samples. Similarly, *TaPR2* also showed significantly lower expression in Δ*Zt-11*-infected wheat leaves at 2 dpi and 6 dpi compared to WT-infected samples. Similar results were reported recently, where deletion strains of *Z. tritici* adenylate cyclase enzyme (Δ*ztcyr1*) were unable to induce activation of the wheat defence response (Child et al., 2023). It was found that Δ*ztcyr1-*infected samples showed down-regulation of *TaPR* genes (Child et al., 2023). Our results that Δ*Zt-11* strains do not activate wheat immune response, potentially due to the absence of recognition via wheat proteins TaSRTRG6, TaSSP6 and TaSSP7 (or guards thereof). There is subsequently no SA accumulation, *PR* gene expression and, as a result, delayed necrosis symptoms compared to WT-infected samples. Reactive oxygen species (ROS) during *Z. tritici-* host interactions are important for resistance to host defences (Shetty et al., 2003, 2007; Reilly et al., 2021). Superoxide dismutase (SOD) and Catalase (CAT) are known ROS scavengers and play a key role in antioxidant defence (Hückelhoven and Kogel, 2003; Nimchuk et al., 2003). However, we did not observe any significant differences in expression of *TaSOD* and *TaCAT* between WT and Δ*Zt-11*-infected wheat samples nor on media supplemented with H_2_O_2_. This suggests no direct role of *Zt-11* in host ROS metabolism and a possible action of Zt11 independent of host ROS production.

## CONCLUSION

In summary, our work characterized a cysteine rich *Z. tritici* specific effector protein Zt-11, for a role in disease. We demonstrated that *Zt-11* is a pathogen secreted protein with high expression at the transition phase during infection. *Zt-11* contributes to pathogen aggressiveness and timely disease progression potentially due to activation of wheat SA responses including *PR* genes.

However, the precise molecular mechanism of how *Zt-11* functions with TaSRTRG6, TaSSP6 and TaSSP7 during the *Z. tritici*-wheat interaction remains to be elucidated.

## Author contributions

SK and AF designed the research. SK and PP carried out all the experiments. PP, CL, NM and AT contributed to generation of deletion mutants and screening. SK and AF wrote the manuscript. JB and AF reviewed the manuscript.

## Acknowledgements

The authors would like to thank Dr A. Bailey, University of Bristol, UK for providing us with the pCAMBIA0380_YA vector and IPO323 strains and Prof. F. Doohan for providing wheat cultivar longbow seeds used in the experiments.

---

**Supplementary Figure 1. Generation of the Δ*Zt11* replacement knock-out mutants**. **A,** Diagram showing the location of the two primer pairs used to confirm successful transformation of *Z. tritici*. The replacement of *Zt11* by the hygromycin-trpC resistance cassette through homologous recombination is depicted by dotted blue lines on the flanking regions. B, Successful disruption of the *Zt11* gene in the mutants indicated by presence of the KO amplicon (2340 base pairs) and absence of the *Zt11* wild-type WT amplicon (2124 base pairs). 1kb (kilobase) ladder (Bioline).

**Supplementary Figure 2. Confirmation of the Δ*Zt11* replacement mutants by amplicon sequencing.** Each row represents sequences (left flank, hygromycin and right flank) confirmed by amplicon sequencing in the mutant strains.

**Supplementary Figure 3. Representative images of pycnidia.** Δ*Zt11* mutant strain, WT (IPO323) and ectopic strain pycnidia on wheat cv. Longbow leaves at 21 dpi. Arrows indicate the cirrhus containing pycnidiospores, scale bars representative of 50µm.

**Supplementary Figure 4. *in vitro* phenotyping of the Δ*Zt11* mutants compared with WT strain.** Approximately 10^7^ spores/mL of spores was spotted on to three different solid growth media; PDA, YPDA and YEG (supplemented with H_2_O_2_ at concentrations 0, 2, 4 and 6mM) and incubated for 7 days at 20°C. Scale bar = 5mm. Images are representative of three independent experiments, with a total of 2 plates per experiment per media type (n=4).

**Supplementary Table 1**. List of Primers used in this study

**Supplementary Table 2.** BlastP of eight *Z. tritici* SSPs previously shown to interact with host wheat proteins TaSSP7, TaSSP6, TaSRTRG6 and TaSRTRG7 (Brennan et al., 2020; Zhou et al., 2020).

