## Supplementary figures and images for "The *Zymoseptoria tritici* effector Zt-11 contributes to aggressiveness in wheat"

### Supplementary fig 1 (1).tiff

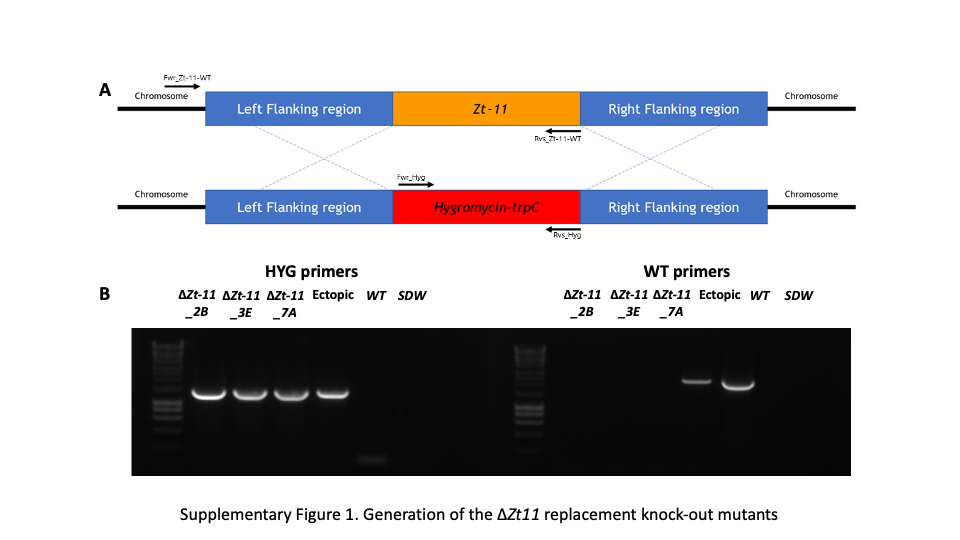

### Supplementary Fig 2 (2).tiff

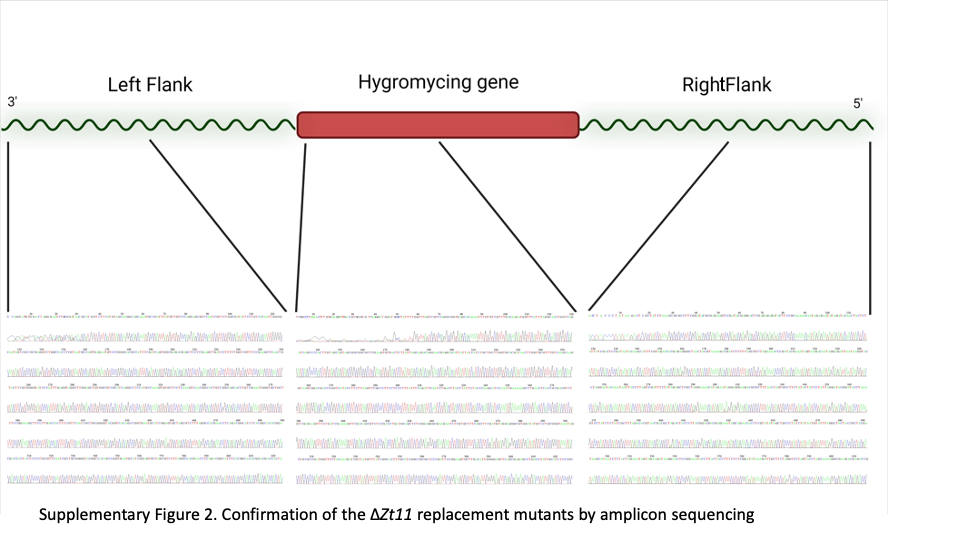

### Supplementary fig 3 (1).tiff

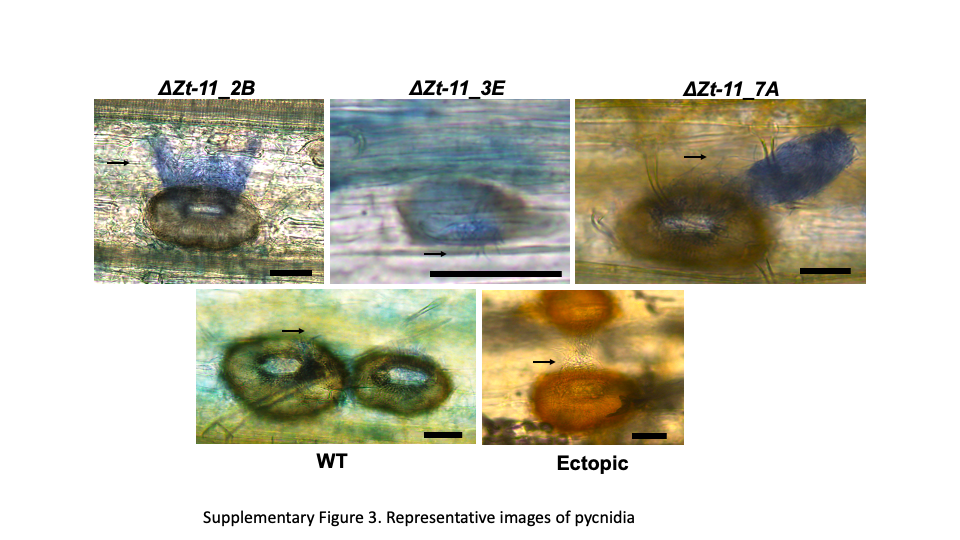

### Supplementary Fig 4 (2).tiff

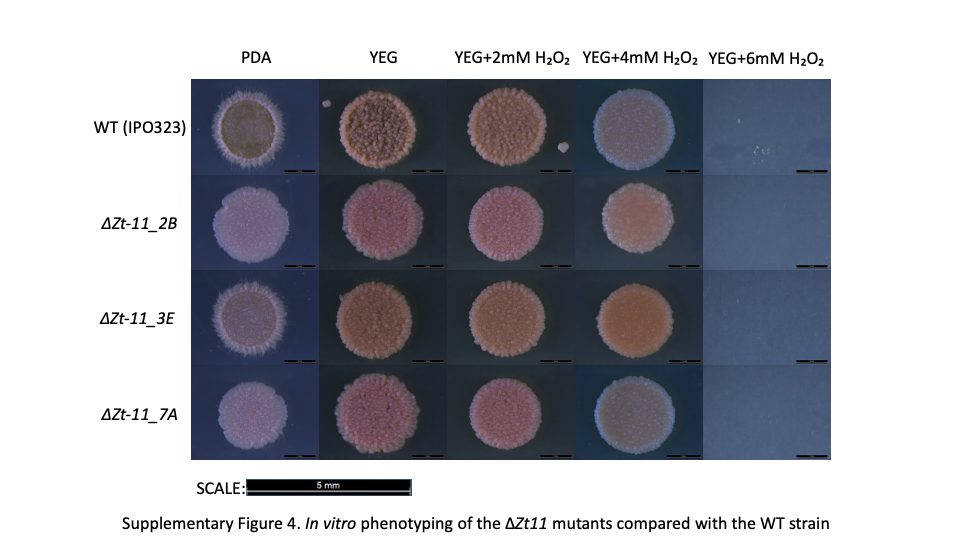
